## Supplemental File 1 for "How do host population dynamics impact Lyme disease risk dynamics in theoretical models?"

1. $E_{wk+1} = \left\{ \begin{array}{l} 0 \text{ if } CDW < CDW_{dev,a} \\ fec \cdot (EA_{wk} + IEA_{wk}) \text{ if } CDW >= CDW_{dev,a} \end{array} \right\} + \left( E_{wk} - \left\{ \begin{array}{l} 0 \text{ if } CDW < CDW_{dev,e} \\ E_{wk} \text{ if } CDW >= CDW_{dev,e} \end{array} \right\} \right) \cdot N(\mu_{t,i}, \sigma_{t,i}) \cdot N(\mu_{p,i}, \sigma_{p,i})$

2. $HL_{wk+1} = \left\{ \begin{array}{l} 0 \text{ if } CDW < CDW_{dev,e} \\ E_{wk} \text{ if } CDW >= CDW_{dev,e} \end{array} \right\} \cdot S_{h,l}$

3. $QL_{wk+1} = (QL_{wk} - \sum_{x=1}^{3}(QL_{wk} \cdot (a_x \cdot H_x)^b) \cdot N(\mu_{a,i}, \sigma_{a,i})) \cdot (N(\mu_{t,i}, \sigma_{t,i}) \cdot N(\mu_{p,i}, \sigma_{p,i}))$

4. $OL_{wk+1} = \sum_{x=1}^{3}(QL_{wk} \cdot (a_x \cdot H_x)^b) \cdot N(\mu_{a,i}, \sigma_{a,i}) \cdot (N(\mu_{t,i}, \sigma_{t,i}) \cdot N(\mu_{p,i}, \sigma_{p,i}))$

5a. $EL_{wk+1} = \left( EL_{wk} - \left\{ \begin{array}{l} 0 \text{ if } CDW < CDW_{dev,l} \\ EL_{wk} \text{ if } CDW >= CDW_{dev,l} \end{array} \right\} \right) \cdot N(\mu_{t,i}, \sigma_{t,i}) \cdot N(\mu_{p,i}, \sigma_{p,i}) + \sum_{x=1}^{3} \left( OL_{wk} \cdot \left\{ \begin{array}{l} S_{x,i} \text{ if } EI_{l,wk} > EI_{max} \\ \frac{S_{m,i} - S_{x,i}}{EI_{max} - EI_{min}} \text{ if } EI_{min} < EI_{l,wk} < EI_{max} \\ S_{m,i} \text{ if } EI_{l,wk} < EI_{min} \end{array} \right\} \cdot C_x (1 - \frac{IH_x}{IH_x + H_x}) \right)$

5b. $IEL_{wk+1} = \left( IEL_{wk} - \left\{ \begin{array}{l} 0 \text{ if } CDW < CDW_{dev,l} \\ IEL_{wk} \text{ if } CDW >= CDW_{dev,l} \end{array} \right\} \right) \cdot N(\mu_{t,i}, \sigma_{t,i}) \cdot N(\mu_{p,i}, \sigma_{p,i}) + \sum_{x=1}^{3} \left( OL_{wk} \cdot \left\{ \begin{array}{l} S_{x,i} \text{ if } EI_{l,wk} > EI_{max} \\ \frac{S_{m,i} - S_{x,i}}{EI_{max} - EI_{min}} \text{ if } EI_{min} < EI_{l,wk} < EI_{max} \\ S_{m,i} \text{ if } EI_{l,wk} < EI_{min} \end{array} \right\} \cdot C_x \frac{IH_x}{IH_x + H_x} \right)$

7a. $HN_{wk+1} = \left\{ \begin{array}{l} 0 \text{ if } CDW < CDW_{dev,n} \\ EL_{wk} \text{ if } CDW >= CDW_{dev,n} \end{array} \right\} \cdot S_{h,n}$

7b. $IHN_{wk+1} = \left\{ \begin{array}{l} 0 \text{ if } CDW < CDW_{dev,n} \\ IEL_{wk} \text{ if } CDW >= CDW_{dev,n} \end{array} \right\} \cdot S_{h,n}$

8a. $QN_{wk+1} = (QN_{wk} - \sum_{x=1}^{3}(QN_{wk} \cdot (a_x \cdot H_x)^b) \cdot N(\mu_{a,i}, \sigma_{a,i})) \cdot (N(\mu_{t,i}, \sigma_{t,i}) \cdot N(\mu_{p,i}, \sigma_{p,i}))$

8b. $IQN_{wk+1} = (IQN_{wk} - \sum_{x=1}^{3}(IQN_{wk} \cdot (a_x \cdot H_x)^b) \cdot N(\mu_{a,i}, \sigma_{a,i})) \cdot (N(\mu_{t,i}, \sigma_{t,i}) \cdot N(\mu_{p,i}, \sigma_{p,i}))$

9a. $ON_{wk+1} = \sum_{x=1}^{3}(QN_{wk} \cdot (a_x \cdot H_x)^b) \cdot N(\mu_{a,i}, \sigma_{a,i}) \cdot (N(\mu_{t,i}, \sigma_{t,i}) \cdot N(\mu_{p,i}, \sigma_{p,i}))$

9b. $ION_{wk+1} = \sum_{x=1}^{3}(IQN_{wk} \cdot (a_x \cdot H_x)^b) \cdot N(\mu_{a,i}, \sigma_{a,i}) \cdot (N(\mu_{t,i}, \sigma_{t,i}) \cdot N(\mu_{p,i}, \sigma_{p,i}))$

10a. $EN_{wk+1} =$

$$\left(EN_{wk} - \begin{cases} 0 \text{ if } CDW < CDW_{dev,n} \\ EN_{wk} \text{ if } CDW >= CDW_{dev,n} \end{cases}\right) \cdot N(\mu_{t,i}, \sigma_{t,i}) \cdot N(\mu_{p,i}, \sigma_{p,i}) +$$

$$\sum_{x=1}^{3} \left( ON_{wk} \cdot \begin{cases} S_{x,i} \text{ if } EL_{n,wk} > EI_{max} \\ \frac{S_{m,i} - S_{x,i}}{EI_{max} - EI_{min}} \text{ if } EI_{min} < EL_{n,wk} < EI_{max} \\ S_{m,i} \text{ if } EL_{n,wk} < EI_{min} \end{cases} \cdot C_x \left(1 - \frac{IH_x}{IH_x + H_x}\right) \right)$$

10a. $IEN_{wk+1} = \left(IEN_{wk} - \begin{cases} 0 \text{ if } CDW < CDW_{dev,n} \\ IEN_{wk} \text{ if } CDW >= CDW_{dev,n} \end{cases}\right) \cdot N(\mu_{t,i}, \sigma_{t,i}) \cdot$

$$N(\mu_{p,i}, \sigma_{p,i}) + \sum_{x=1}^{3} \left( ION_{wk} \cdot \begin{cases} S_{x,i} \text{ if } EL_{n,wk} > EI_{max} \\ \frac{S_{m,i} - S_{x,i}}{EI_{max} - EI_{min}} \text{ if } EI_{min} < EL_{n,wk} < EI_{max} \\ S_{m,i} \text{ if } EL_{n,wk} < EI_{min} \end{cases} \right) +$$

$$\sum_{x=1}^{3} \left( ON_{wk} \cdot \begin{cases} S_{x,i} \text{ if } EL_{n,wk} > EI_{max} \\ \frac{S_{m,i} - S_{x,i}}{EI_{max} - EI_{min}} \text{ if } EI_{min} < EL_{n,wk} < EI_{max} \\ S_{m,i} \text{ if } EL_{n,wk} < EI_{min} \end{cases} \cdot C_x \left(\frac{IH_x}{IH_x + H_x}\right) \right)$$

11a. $HA_{wk+1} = \begin{cases} 0 \text{ if } CDW < CDW_{dev,a} \\ EN_{wk} \text{ if } CDW >= CDW_{dev,a} \end{cases} \cdot S_{h,a}$

11b. $IHA_{wk+1} = \begin{cases} 0 \text{ if } CDW < CDW_{dev,a} \\ IEA_{wk} \text{ if } CDW >= CDW_{dev,a} \end{cases} \cdot S_{h,a}$

12a. $QA_{wk+1} =$

$$(QA_{wk} - \sum_{x=1}^{3} (QA_{wk} \cdot (a_x \cdot H_x)^b) \cdot N(\mu_{a,a}, \sigma_{a,a})) \cdot (N(\mu_{t,a}, \sigma_{t,a}) \cdot N(\mu_{p,a}, \sigma_{p,a}))$$

12b. $IQA_{wk+1} =$

$$(IQA_{wk} - \sum_{x=1}^{3} (IQA_{wk} \cdot (a_x \cdot H_x)^b) \cdot N(\mu_{a,a}, \sigma_{a,a})) \cdot (N(\mu_{t,a}, \sigma_{t,a}) \cdot N(\mu_{p,a}, \sigma_{p,a}))$$

13a. $OA_{wk+1} = \sum_{x=1}^{3} (QA_{wk} \cdot (a_x \cdot H_x)^b) \cdot N(\mu_{a,a}, \sigma_{a,a}) \cdot (N(\mu_{t,a}, \sigma_{t,a}) \cdot N(\mu_{p,a}, \sigma_{p,a}))$

13b. $IOA_{wk+1} = \sum_{x=1}^{3} (IQA_{wk} \cdot (a_x \cdot H_x)^b) \cdot N(\mu_{a,a}, \sigma_{a,a}) \cdot (N(\mu_{t,a}, \sigma_{t,a}) \cdot N(\mu_{p,a}, \sigma_{p,a}))$

14a. $EA_{wk+1} =$

$$\left(EA_{wk} - \begin{cases} 0 \text{ if } CDW < CDW_{dev,a} \\ EN_{wk} \text{ if } CDW >= CDW_{dev,a} \end{cases}\right) \cdot N(\mu_{t,a}, \sigma_{t,a}) \cdot N(\mu_{p,a}, \sigma_{p,a}) +$$

$$\sum_{x=1}^{3} \left( OA_{wk} \cdot \begin{cases} S_{x,a} \text{ if } EL_{a,wk} > EI_{max} \\ \frac{S_{m,a} - S_{x,a}}{EI_{max} - EI_{min}} \text{ if } EI_{min} < EL_{a,wk} < EI_{max} \\ S_{m,a} \text{ if } EL_{a,wk} < EI_{min} \end{cases} \cdot C_x \left(1 - \frac{IH_x}{IH_x + H_x}\right) \right)$$

14a. $$IEA_{wk+1} = \left( IEA_{wk} - \left\{ \begin{array}{l} 0 \text{ if } CDW < CDW_{dev,a} \\ IEA_{wk} \text{ if } CDW >= CDW_{dev,a} \end{array} \right\} \right) \cdot N(\mu_{t,a}, \sigma_{t,a}) \cdot N(\mu_{p,a}, \sigma_{p,a}) + \sum_{x=1}^{3} \left( ION_{wk} \cdot \left\{ \begin{array}{l} S_{x,a} \text{ if } EL_{a,wk} > EI_{max} \\ \frac{S_{m,a} - S_{x,a}}{EI_{max} - EI_{min}} \text{ if } EI_{min} < EL_{a,wk} < EI_{max} \\ S_{m,a} \text{ if } EL_{a,wk} < EI_{min} \end{array} \right\} \right) + \sum_{x=1}^{3} \left( OA_{wk} \cdot \left\{ \begin{array}{l} S_{x,a} \text{ if } EL_{a,wk} > EI_{max} \\ \frac{S_{m,a} - S_{x,a}}{EI_{max} - EI_{min}} \text{ if } EI_{min} < EL_{a,wk} < EI_{max} \\ S_{m,a}, EL_{a,wk} < EI_{min} \end{array} \right\} \cdot C_x\left(\frac{IH_x}{IH_x + H_x}\right) \right)$$

15a. $$EI_{l,wk} = sum_{k=wk}^{wk-9}(0.44^{(k-1)} \cdot sum_{x=1}^{3} \frac{EI_{scl} \cdot (OL_k + IOL_k)}{H_x})$$

16a. $$SM_{wk+1} = \left\{ \begin{array}{l} SM_{wk} \cdot S_x + (SM_{wk} + IM_{wk}) \cdot B_x, wk \ \% \ 52 \neq 0 \\ \left\{ \begin{array}{l} SM_{wk} + (SM_{wk} - N(\mu_m, \sigma_m)), SM_{wk} - N(\mu_m, \sigma_m) > 0 \\ SM_{wk} + (SM_{wk} - N(\mu_m, \sigma_m)) \cdot \frac{SM_{wk}}{SM_{wk} + IM_{wk}}, SM_{wk} - N(\mu_m, \sigma_m) < 0 \end{array} \right. \end{array} \right.$$

16b. $$IM_{wk+1} = \left\{ \begin{array}{l} IM_{wk} \cdot S_x + T(ITH) \cdot SM_{wk}, wk \ \% \ 52 \neq 0 \\ \left\{ \begin{array}{l} IM_{wk}, SM_{wk} - N(\mu_m, \sigma_m) > 0 \\ SM_{wk} + (SM_{wk} - N(\mu_m, \sigma_m)) \cdot \frac{IM_{wk}}{SM_{wk} + IM_{wk}}, SM_{wk} - N(\mu_m, \sigma_m) < 0 \end{array} \right. \end{array} \right.$$

16c. $$SMM_{wk+1} = SMM_{wk} \cdot S_x + (SMM_{wk} + IMM_{wk}) \cdot B_x$$

16d. $$IMM_{wk+1} = IMM_{wk} \cdot S_x + T(ITH_x) \cdot SMM_{wk}$$

16e. $$SD_{wk+1} = SD_{wk} \cdot S_x + (SD_{wk} + ID_{wk}) \cdot B_x$$

16f. $$ID_{wk+1} = ID_{wk} \cdot S_x + T(ITH_x) \cdot SD_{wk}$$

17. $$ITH_x = TIF \cdot \frac{\sum(IO(L,N,A))}{H_{x,wk} + IH_{x,wk}}$$

**Note**: $T$ is the effective transmission rate, defined in "Infection".
