## Supplementary figures and images for "How do host population dynamics impact Lyme disease risk dynamics in theoretical models?"

### Supplemental Figure 1

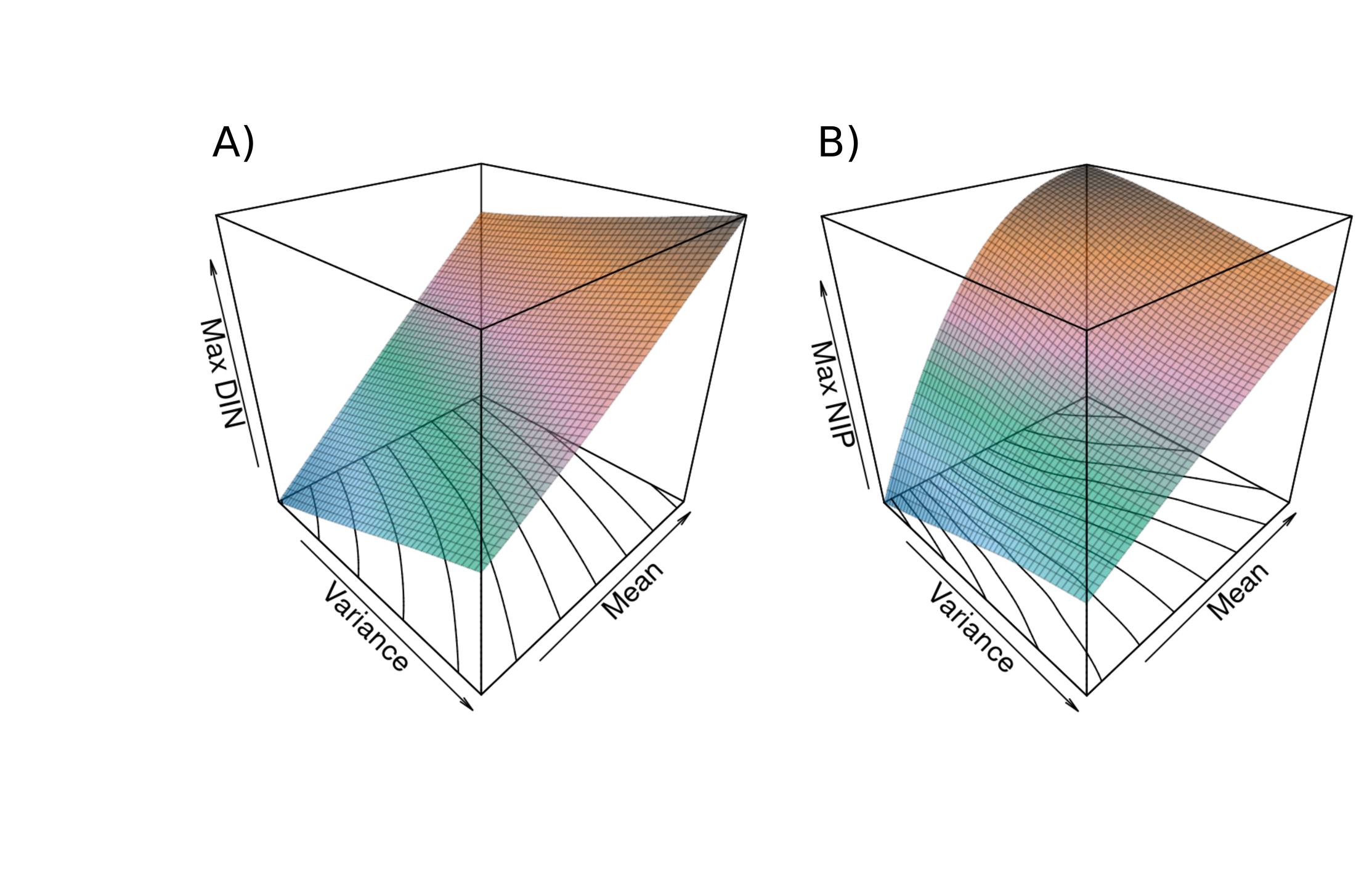

### Supplemental Figure 2

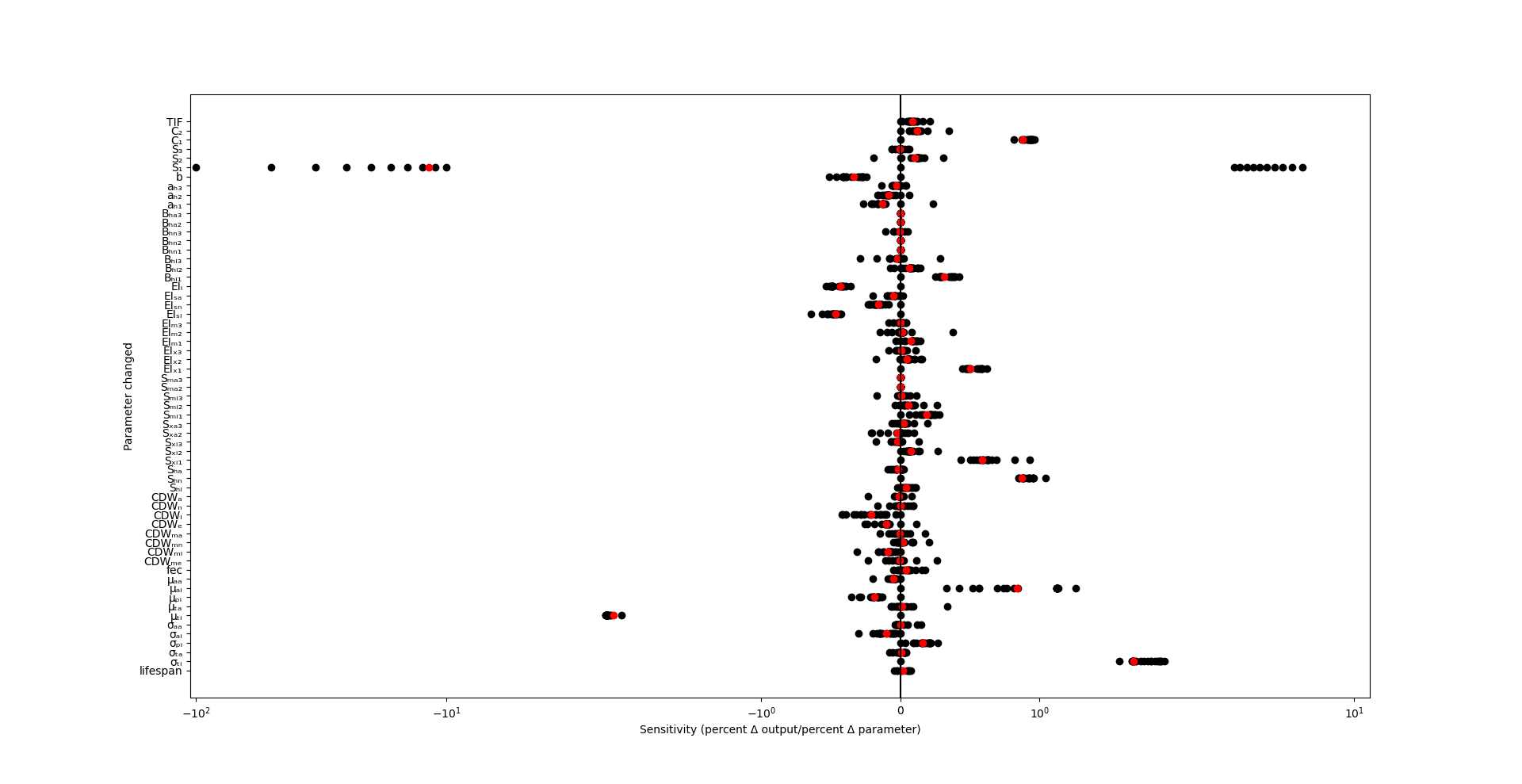
