## Supplemental Table 1 for "How do host population dynamics impact Lyme disease risk dynamics in theoretical models?"

| Disease Metric | variance | min | Q1 | mean | median | Q3 | max |
| --- | --- | --- | --- | --- | --- | --- | --- |
| mean DIN | 0 | 570 | 2031 | 3351 | 3394 | 4694 | 5968 |
| | 1-33 | 579 | 1927 | 3285 | 3270 | 4588 | 7055 |
| | 34-66 | 663 | 2099 | 3280 | 3132 | 4327 | 8047 |
| | 67-99 | 694 | 2412 | 3518 | 3327 | 4495 | 8958 |
| min DIN | 0 | 34 | 125 | 207 | 210 | 291 | 367 |
| | 1-33 | 34 | 121 | 205 | 204 | 286 | 437 |
| | 34-66 | 37 | 131 | 205 | 197 | 273 | 505 |
| | 67-99 | 39 | 147 | 217 | 204 | 281 | 639 |
| max DIN | 0 | 1829 | 6447 | 10677 | 10805 | 14981 | 18980 |
| | 1-33 | 1869 | 7120 | 11304 | 11400 | 15442 | 24159 |
| | 34-66 | 2438 | 9252 | 13122 | 12995 | 16813 | 28904 |
| | 67-99 | 2715 | 11362 | 15319 | 15109 | 19070 | 33706 |
| amp DIN | 0 | 1796 | 6322 | 10470 | 10595 | 14690 | 18613 |
| | 1-33 | 1835 | 7007 | 11099 | 11193 | 15145 | 23760 |
| | 34-66 | 2397 | 9114 | 12917 | 12804 | 16557 | 28412 |
| | 67-99 | 2676 | 11185 | 15102 | 14896 | 18804 | 33067 |
| mean DON | 0 | 1790 | 4101 | 6075 | 6173 | 8109 | 9997 |
| | 1-33 | 1809 | 4110 | 6143 | 6168 | 8099 | 11793 |
| | 34-66 | 2040 | 4717 | 6466 | 6308 | 8092 | 13695 |
| | 67-99 | 2115 | 5441 | 7113 | 6882 | 8693 | 15296 |
| min DON | 0 | 185 | 356 | 506 | 513 | 660 | 800 |
| | 1-33 | 186 | 361 | 513 | 514 | 662 | 956 |
| | 34-66 | 195 | 397 | 534 | 525 | 661 | 1094 |
| | 67-99 | 200 | 441 | 575 | 556 | 699 | 1384 |
| max DON | 0 | 9625 | 18629 | 26569 | 26923 | 34696 | 42160 |
| | 1-33 | 9711 | 21187 | 28779 | 28994 | 36479 | 55245 |
| | 34-66 | 11298 | 27541 | 34850 | 34835 | 41885 | 69202 |
| | 67-99 | 12451 | 33688 | 41572 | 41497 | 49346 | 76826 |
| amp DON | 0 | 5618 | 12297 | 18258 | 18495 | 24348 | 29964 |
| | 1-33 | 5680 | 13962 | 19648 | 19768 | 25348 | 38853 |
| | 34-66 | 6746 | 18062 | 23510 | 23463 | 28733 | 46425 |
| | 67-99 | 7450 | 22130 | 27853 | 27758 | 33435 | 54137 |
| mean NIP | 0 | 0.32 | 0.49 | 0.53 | 0.55 | 0.58 | 0.6 |
| | 1-33 | 0.32 | 0.46 | 0.51 | 0.53 | 0.57 | 0.6 |
| | 34-66 | 0.32 | 0.43 | 0.48 | 0.48 | 0.53 | 0.61 |
| | 67-99 | 0.33 | 0.42 | 0.46 | 0.46 | 0.5 | 0.6 |
| min NIP | 0 | 0.3 | 0.36 | 0.38 | 0.38 | 0.4 | 0.41 |
| | 1-33 | 0.27 | 0.32 | 0.35 | 0.36 | 0.38 | 0.41 |
| | 34-66 | 0.24 | 0.29 | 0.32 | 0.32 | 0.34 | 0.41 |
| | 67-99 | 0.22 | 0.28 | 0.3 | 0.3 | 0.32 | 0.4 |
| max NIP | 0 | 0.32 | 0.53 | 0.57 | 0.6 | 0.63 | 0.65 |
| | 1-33 | 0.32 | 0.52 | 0.56 | 0.59 | 0.62 | 0.66 |
| | 34-66 | 0.34 | 0.5 | 0.55 | 0.56 | 0.6 | 0.66 |
| | 67-99 | 0.35 | 0.5 | 0.54 | 0.54 | 0.59 | 0.66 |
| amp NIP | 0 | 0.03 | 0.17 | 0.19 | 0.21 | 0.23 | 0.24 |
| | 1-33 | 0.03 | 0.19 | 0.21 | 0.23 | 0.24 | 0.28 |
| | 34-66 | 0.05 | 0.21 | 0.23 | 0.24 | 0.26 | 0.31 |
| | 67-99 | 0.07 | 0.21 | 0.24 | 0.24 | 0.27 | 0.33 |
