## Supplemental Table 2 for "How do host population dynamics impact Lyme disease risk dynamics in theoretical models?"

| stage class | description | duration |
|---|---|---|
| Eggs | Beginning (1st stage) of tick development | 52 weeks |
| Hardening larvae | Newly hatched larval (2nd stage) ticks, which must harden | 1 week |
| Questing larvae | Larvae which have not fed, and are able to quest for hosts | 52 weeks |
| On-host larvae | Larvae which have found, and are currently feeding from, a host | 52 weeks |
| Engorged larvae | Larvae which have fed and now must develop into nymphs | 52 weeks |
| Hardening nymphs | Newly hatched nymphal (3nd stage) ticks, which must harden | 1 week |
| Questing nymphs | Nymphs which have not fed, and are able to quest for hosts | 52 weeks |
| On-host nymphs | Nymphs which have found, and are currently feeding from, a host | 52 weeks |
| Engorged nymphs | Nymphs which have fed and now must develop into nymphs | 52 weeks |
| Hardening adults | Newly hatched adult (4th stage) ticks, which must harden | 1 week |
| Questing adults | Adults which have not fed, and are able to quest for hosts | 52 weeks |
| On-host adults | Adults which have found, and are currently feeding from, a host | 52 weeks |
| Engorged adults | Adults which have fed and now must develop into nymphs | 52 weeks |
