## Supplemental Table 3 for "How do host population dynamics impact Lyme disease risk dynamics in theoretical models?"

| stage category | processes |
| --- | --- |
| Egg | survival, development |
| Free living unfed | survival, questing |
| Free living fed | survival, development |
| Hardening | survival |
| On-host | survival, infection, transmission |
