## Supplemental Table 4 for "How do host population dynamics impact Lyme disease risk dynamics in theoretical models?"

| Parameter name | Description | Value | Citation |
|---|---|---|---|
| Tick Lifespan | Maximum lifespan of a tick in egg or free living stages | 52 weeks | * |
| $\sigma_{t,i}$ | Standard deviation of temperature induced survival in larvae & nymphs | 45 | * |
| $\sigma_{t,a}$ | Standard deviation of temperature induced survival in adults | 50 | * |
| $\sigma_{p,i}$ | Standard deviation of precipitation induced survival in all tick stages | 70 | * |
| $\sigma_{a,i}$ | Standard deviation of temperature induced activity in larvae & nymphs | 5 | [1, 2] |
| $\sigma_{a,a}$ | Standard deviation of temperature induced activity in adults | 8 | [1, 2] |
| $\mu_{t,i}$ | Mean of temperature induced survival distribution in larvae & nymphs | 25 | [3–5] |
| $\mu_{t,a}$ | Mean of temperature induced survival distribution in adults | 8 | [3–5] |
| $\mu_{p,i}$ | Mean of precipitation induced survival distribution in all tick stages | 8 | [3–5] |
| $\mu_{a,i}$ | Mean of temperature induced activity distribution in larvae & nymphs | 25 | [1, 2] |
| $\mu_{a,a}$ | Mean of temperature induced activity distribution in adults | 8 | [1, 2] |
| $weeks_s$ | Number of simulation weeks | 2600 weeks | * |
| $Egg_i$ | Initial egg density | 289748 | ** |
| $EL_i$ | Initial engorged larvae density | 1179 | ** |
| $EN_i$ | Initial engorged nymph density | 51 | ** |
| $EA_i$ | Initial engorged adult density | 57 | ** |
| $IEL_i$ | Initial infected engorged larvae density | 1563 | ** |
| $IEN_i$ | Initial infected engorged nymph density | 128 | ** |
| $IEA_i$ | Initial infected engorged adult density | 158 | ** |
| $QL_i$ | Initial questing larvae density | 312150 | ** |
| $QN_i$ | Initial questing nymph density | 2609 | ** |
| $QA_i$ | Initial questing adult density | 121 | ** |
| $IQN_i$ | Initial infected questing nymph density | 2897 | ** |
| $IQA_i$ | Initial infected questing adult density | 257 | ** |
| $IH_i$ | Initial hardening ticks (all stages) | 0 ind. | ** |
| $OH_i$ | Initial ticks on hosts (all stages) | 0 ind. | ** |
| $\sigma_m$ | Standard deviation of mouse density | 0-99 | * |
| $\mu_m$ | Mean mouse density | 0-99 | * |
| fec | fecundity | 860 | [6] |
| $CDW_{min}$ | Minimum degree week for development by CDW (all stages) | 6 | [5] |
| $CDW_{dev,e}$ | Threshold CDW for development of eggs | 110 | [5] |
| $CDW_{dev,l}$ | Threshold CDW for development of larvae | 58 | [5] |
| $CDW_{dev,n}$ | Threshold CDW for development of nymphs | 81 | [5] |
| $CDW_{dev,a}$ | Threshold CDW for egg laying by adults | 28 | [5] |
| $S_{h,l}$ | Hardening larvae survival | 0.9647684 | [4] |
| $S_{h,n}$ | Hardening nymph survival | 0.9994695 | [4] |
| $S_{h,a}$ | Hardening adult survival | 0.9999692 | [4] |

* values chosen as part of study.

** Initial conditions taken from week 0 of the last year of a simulation with mean mouse density set to 40 and no variation.
