## Supplemental Table 5 for "How do host population dynamics impact Lyme disease risk dynamics in theoretical models?"

| Parameter name | Description | Value | Citation |
|---|---|---|---|
| $S_{x,i}$ | Maximum on-host immature survival for each host (mice, medium, deer) | 0.6 | [1] |
| $S_{x,a}$ | Maximum on-host adult survival for each host (mice, medium, deer) | (0, 0.6, 0.7746) | [1] |
| $S_{m,i}$ | Minimum on-host immature survival for each host (mice, medium, deer) | 0.2 | [1] |
| $S_{m,a}$ | Minimum on-host adult survival for each host (mice, medium, deer) | (0, 0.2, 0.4474) | [1] |
| $EI_{max}$ | Maximum engorgement index for each host (mice, medium, deer), and all on host stages | (0.75, 4, 300) | [1] |
| $EI_{min}$ | Minimum engorgement index for each host (mice, medium, deer), and all on host stages | (0.15, 0.8, 60) | [1] |
| $EI_{scl}$ | Engorgement index allometric scaling coefficient for (larval, nymphal, adult) tick density | (0.0021, 0.014, 1) | [1] |
| $EI_{loss}$ | Engorgement index weekly loss | 0.44 | [1] |
| $B_{h,l}$ | Host larval burdens for each host (mice, medium, deer) | (100, 200, 1000) | [1] |
| $B_{h,n}$ | Host nymphal burdens for each host (mice, medium, deer) | (20, 100, 500) | [1] |
| $B_{h,a}$ | Host adult burdens for each host (mice, medium, deer) | (0, 20, 100) | [1] |
| $a_h$ | Basal host finding rate for each host (mice, medium, deer), and all tick stages | (0.01, 0.025, 0.05) | [1] |
| $b$ | Host finding rate scaling for all tick stages and species | 0.515 | [1] |
| $S_{mice}$ | Mouse survival | 0.9808 | * |
| $S_{med}$ | Medium host survival | 0.9904 | * |
| $S_{deer}$ | Deer survival | 0.9936 | * |
| $C_{mice}$ | Mouse competency | 0.75 | [2, 3] |
| $C_{med}$ | Medium host competency | 0.5 | [3–5] |
| $C_{deer}$ | Deer competency | 0 | [4, 6, 7] |
| $TIF$ | Tick infectivity factor, tick-to-host transmission | 0.9 | [1] |

* values chosen as part of study.

** Initial conditions taken from week 0 of the last year of a simulation with mean mouse density set to 40 and no variation.
